# Herpes Simplex Virus-1 targets the 2’-3’cGAMP importer SLC19A1 as an antiviral countermeasure

**DOI:** 10.1101/2024.01.24.577105

**Authors:** Zsuzsa K. Szemere, Eain A. Murphy

**Affiliations:** Microbiology and Immunology Department, SUNY-Upstate Medical University, Syracuse NY 13210

## Abstract

To establish a successful infection, herpes simplex virus-1 (HSV-1), a virus with high seropositivity in the human population, must undermine host innate and intrinsic immune defense mechanisms, including the stimulator of interferon genes (STING) pathway. Recently it was discovered that not only *de novo* produced intracellular 2’-3’cGAMP, but also extracellular 2’-3’cGAMP activates the STING pathway by being transported across the cell membrane via the folate transporter, SLC19A1, the first identified extracellular antiporter of this signaling molecule. We hypothesized that the import of exogenous 2’-3’cGAMP functions to establish an antiviral state like that seen with the paracrine antiviral activities of interferon. Further, to establish a successful infection, HSV-1 must undermine this induction of the STING pathway by inhibiting the biological functions of SLC19A1. Herein, we report that treatment of the monocytic cell line, THP-1 cells, epithelial cells (ARPE-19) and SH-SY5Y neuronal cell line with exogenous 2’-3’cGAMP induces interferon production and establishes an antiviral state. Using either pharmaceutical inhibition or genetic knockout of SLC19A1 blocks the 2’-3’cGAMP-induced antiviral state. Additionally, HSV-1 infection results in the reduction of SLC19A1 transcription, translation, and importantly, the rapid removal of SLC19A1 from the cell surface of infected cells. Our data indicate SLC19A1 functions as a newly identified antiviral mediator for extracellular 2’-3’cGAMP which is undermined by HSV-1. This work presents novel and important findings about how HSV-1 manipulates the host’s immune environment for viral replication and discovers details about an antiviral mechanism which information could aid in the development of better antiviral drugs in the future.

**Importance:** HSV-1 has evolved multiple mechanisms to neutralize of the host’s innate and intrinsic defense pathways, such as the STING pathway. Here, we identified an antiviral response in which extracellular 2’-3’cGAMP triggers IFN production via its transporter SLC19A1. Moreover, we report that HSV-1 blocks the functions of this transporter thereby impeding the antiviral response, suggesting exogenous 2’-3’cGAMP can act as an immunomodulatory molecule in uninfected cells to activate the STING pathway, and priming an antiviral state, similar to that seen in interferon responses. The details of this mechanism highlight important details about HSV-1 infections. This work presents novel findings about how HSV-1 manipulates the host’s immune environment for viral replication and reveals details about a novel antiviral mechanism. These findings expand our understanding of how viral infections undermine host responses and may help in the development of better broad based antiviral drugs in the future.

## INTRODUCTION

The stimulator of interferon genes (STING) pathway is a pivotal component of the innate immune system. The central biological function of STING upon binding cyclic dinucleotides (cDNs) is to post translationally modify host transcriptional activators, resulting in the production of type I IFNs and cytokines, thus making it an essential antiviral defense mechanism (1–3). The genomes of DNA viruses, and to a lesser degree, RNA viruses, are activators of the STING pathway (4,5). In the case of DNA viruses, such as the herpesvirus family, cytosolic DNA is recognized by cyclic GMP-AMP synthase (cGAS) to produce 2′3′-cyclic GMP-AMP (2′3′-cGAMP) from ATP and GTP. Subsequent binding of 2′3′-cGAMP to STING results in autophosphorylation of STING followed by translocation to the ER to Golgi intermediate compartment (ERGIC). Here, it recruits and activates TANK binding kinase 1 (TBK1), which phosphorylates the transcription factors interferon regulatory factor 3 (IRF3) and the IKK complex, amongst other substrates. Upon phosphorylation, IRF3 dimerizes, enters the nucleus, and activates the promoter of type I interferon. TBK1 phosphorylation of IKK releases cytoplasmic retention of nuclear factor κ-light chain enhancer of activated B cells (NF-κB), resulting in its translocation to the nucleus and induction of NF-κB-dependent cytokine transcription (6–8).

Accordingly, successful DNA and RNA viruses have developed mechanisms to undermine the STING pathway in order to bolster their replication. Herpes simplex virus-1 (HSV-1), a DNA virus that has co-evolved with its host, employs various strategies to evade the host innate and intrinsic immune defense mechanisms, including, but not limited to, disruption of the STING antiviral pathway at multiple junctures. For instance, HSV-1 interferes with the biological functions of cGAS, STING, and downstream signaling molecules, like TBK1 and IRF3 (9). For example, the viral protein, ICP0, plays a central role in this inhibition by binding to and inhibiting STING, though the extent of its functions remains undefined (10). Additionally, the immediate early (IE) HSV-1 protein, ICP27, undermines the STING pathway by binding to and inhibiting activated TBK1 (11). Thus, while HSV-1 exhibits redundancy in its inhibition of the STING pathway, our understanding of HSV-1’s ability to block antiviral responses remains incomplete.

As highlighted above, 2’-3’cGAMP, the secondary messenger molecule, is key in activating the STING pathway, leading to the production of IFNs (12). Historically, 2’-3’cGAMP synthesized *de novo* by cGAS was viewed primarily as an intracellular messenger. However, recent studies have explored its role as an extracellular mediator that impacts cells in the absence of activated cGAS. (13–15) The addition of extracellular 2’-3’cGAMP was assessed for its potential to boost the host’s immune response against tumors that are dependent on STING pathway activation(16). Cancer cells frequently release 2’-3’cGAMP into the extracellular space, where it is hydrolyzed by ectonucleotide pyrophosphatase/phosphodiesterase 1 (ENPP1). Blocking ENPP1 increased extracellular 2’-3’cGAMP levels, which subsequently delayed tumor growth (17). Additional research revealed 2’-3’cGAMP is transferred between cells through gap junctions or packaged into viral particles, in turn, effectively transmitting the immune response to neighboring cells.(18,19) This underscores the significance of both intracellular and extracellular 2’-3’cGAMP in activating the STING pathway. Given its central role in innate immune responses, studying 2’-3’cGAMP’s functional mechanisms as an immunotransmitter during disease states is vital for antiviral and anticancer therapeutic advancements.

Due to the limited membrane permeability of cDNs, the means by which extracellular 2’-3’cGAMP crosses the cell membrane remained unclear until recently. Solute carrier family 19 member 1 (SLC19A1; a.k.a. Reduced Folate Carrier 1 (RFC1)) was identified in a CRISPR knockout screen as a functional transporter of exogenous 2’-3’cGAMP (14,20). Originally thought to only transport reduced folates and antifolates across the cell membrane (21), SLC19A1 effectively transports extracellular 2’-3’cGAMP, allowing it to then bind and active the STING pathway in a non-canonical, cGAS-independent fashion (14).Moreover, other investigated members of the SLC family of folate transporters demonstrate little or no specificity in the uptake of cDNs (14). In light of this, research on exogenous 2’-3’cGAMP treatment is gaining traction due to its therapeutic potential in cancer treatment and its emerging role in intracellular communication.

Currently the link between extracellular 2’-3’cGAMP and viral infections remains undefined. Based on the pattern of a non-canonical cGAS-independent STING activation in cancer cells, we hypothesize that extracellular 2’-3’cGAMP leads to an antiviral state by activating the STING pathway in THP-1 and ARPE-19 cells that is mediated by SLC19A1 antiporter. Considering HSV-1’s capacity to inhibit the STING pathway at multiple stages (22–24), we further hypothesize that SLC19A1 is a target for viral regulation. Herein, we show that exogenous treatment with 2’-3’cGAMP induces hallmarks of an antiviral state within cell types of distinct origin. Further, pre-treatment of cells with exogenous 2’-3’cGAMP limits HSV-1 replication, and this is dependent on the expression of SLC19A1, as either pharmacological inhibition or genetic disruption of this protein abrogates the antiviral effects of 2’-3’cGAMP treatment. Importantly, the virus rapidly removes SLC19A1 cell surface expression upon infection. Finally, infection by HSV-1 reduces the RNA, total protein levels and cell surface expression of the recently identified transporter, SLC19A1. These data reveal extracellular 2-3’cGAMP, imported by SLC19A1 establishing an antiviral state, which HSV-1 undermines.

## RESULTS

### Exogenous treatment with 2’-3’cGAMP establishes an antiviral state *in vitro*

SLC19A1 import of extracellular 2’-3’cGAMP induces the STING pathway in a cGAS-independent mechanism in cancer cells (14). Thus, we wanted to determine if extracellular 2’-3’cGAMP uptake in cells permissive for HSV-1 initiates an antiviral transcriptional profile, similar to that seen with canonical cGAS-generated 2’-3’cGAMP. While a majority of studies with HSV-1 utilize Vero cells derived from the kidney of an African green monkey, this cell line has a defective interferon response(25,26). Thus, we elected to utilize cells of human origin. To this end, we treated THP-1 and ARPE-19 cells with increasing concentrations of extracellular 2’-3’cGAMP and monitored the impact on cell viability and found exogenous 2’-3’cGAMP was well-tolerated in each cell type at the assessed concentrations and duration (Fig 1A). To determine the impact of adding 2’-3’cGAMP to the media, cells were treated with the cyclic dinucleotide for 24 hrs and quantified *IFNB* RNA levels by RT-qPCR as a surrogate readout for the activation of the STING pathway (27). In both THP-1 and ARPE-19 cells, there was a dose-dependent increase of *IFNB* RNA expression in response to treatment (Fig 1B), indicating the cells responded to exogenous 2’-3’cGAMP. Compared to THP-1 cells, ARPE-19 cells showed a less prominent *IFNB* transcriptional activation. This reduced responsiveness could be explained by the potential difference in the level of SLC19A1 expression or the degree of STING stimulation exhibited by different cell types. Overall, these data suggest extracellular 2’-3’cGAMP treatment of cells initiates an antiviral, IFN-producing state.

**Figure 1.**
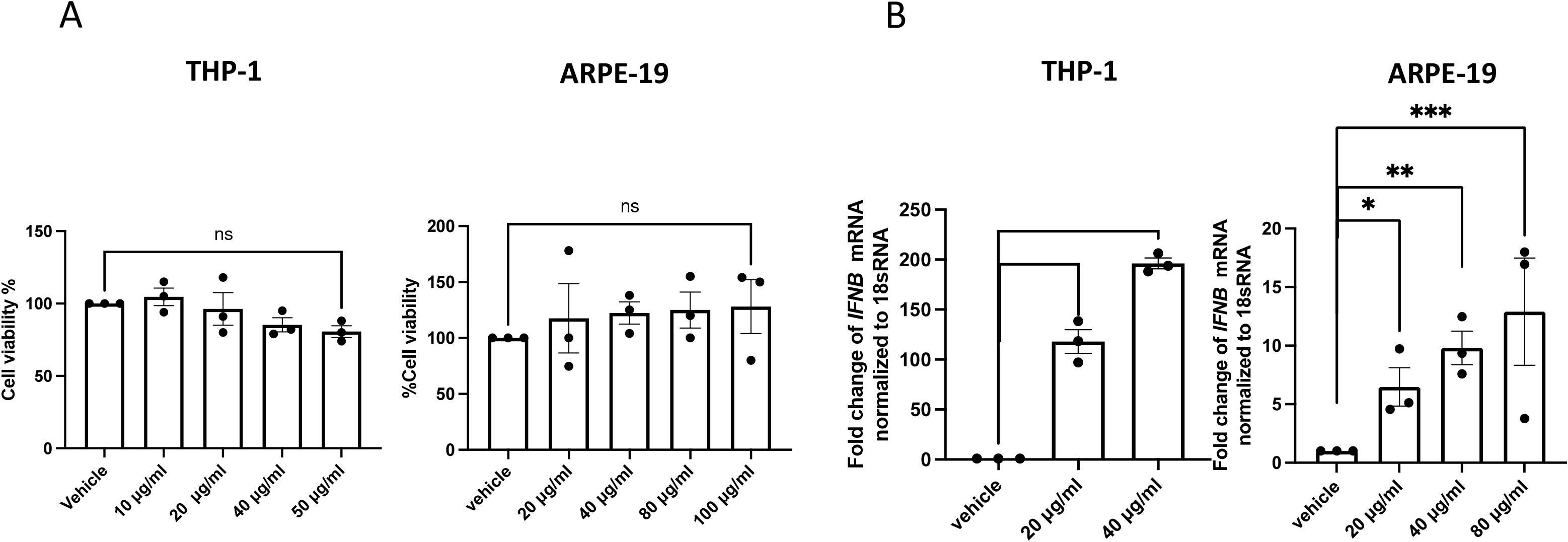
Exogenous 2’-3’cGAMP induces an IFN response in THP-1 and ARPE-19 cells. (A) THP-1 and ARPE-19 cells were treated with increasing concentrations of 2’-3’cGAMP for 24 h and cell viability after the treatment with the drug was assessed by MTT assay. (B) The fold change of *IFNB* mRNA levels in THP-1 and ARPE-19 cells in response to increasing concentration of 2’-3’cGAMP for 24 h were measured by RT-QPCR and normalized to cellular *18sRNA*. n=3. ****= pValue <0.0001, ***=0.001, **=0.01, *=0.05.

It is well-accepted that paracrine interferon treatment can establish an antiviral state within uninfected cells. Therefore, we next asked whether pre-treatment with extracellular 2’-3’cGAMP was sufficient to establish a cellular environment that affects viral replication, similar to interferon treatment. To this end, we first evaluated HSV-1 susceptibility and permissiveness of the cell types used in our study. We infected Vero, THP-1 or ARPE-19 cells with HSV-1 US11eGFP Patton strain (28) and confirmed that each cell type is susceptible and permissive for HSV-1. Importantly, we found each cell type supported robust viral replication (Fig 2A). To determine the effects of pretreatment with extracellular 2’-3’cGAMP on total viral yield, we measured cell-free viral titers in the presence and absence of the CDN. Pre-treatment of cells with exogenous 2’-3’cGAMP resulted in a reduction in viral titers in THP-1 cells, however we did not observe this attenuation in pretreated ARPE-19 cells (Fig 2B), consistent with the significantly reduced induction of *IFNB* (>1 log) in the ARPE-19 cells when compared to comparable treated THP-1 cells (Fig 1B). Next, as HSV-1 is a neurotropic virus, we wanted to test the impact of exogenous 2’-3’cGAMP treatment on cells of neuronal origin. We utilized the neuroblastoma derived cell line, SH-SY5Y, which is a permissive human neuronal cell line capable of productive HSV-1 infection (Fig 2B) (29). We found that similar to the results in THP-1 cells, pre-treatment with extracellular 2’-3’cGAMP resulted in a decreased viral yield in SH-Sy5Y cells as well (Fig 2B). To determine the stage at which HSV-1 is affected by 2’-3’cGAMP treatment, THP-1 cells were infected with a HSV-1 strain that expresses tdTomato under IE kinetics(30), thereby allowing us to monitor events upon viral infection. We focused on THP-1 cells for this experiment since they are the most responsive to extracellular 2’-3’cGAMP treatment (Fig 1B). THP-1 cells were pre-treated as above and infected with tdTomato virus. We then measured IE-expressed tdTomato gene expression by RT-QPCR. Compared to vehicle treated, infected cells, 2’-3’cGAMP pre-treated cells showed a significant decrease in IE gene reporter expression in THP-1 cells (Fig 2C), suggesting extracellular 2’-3’cGAMP mediates an antiviral state that inhibits HSV-1 replication at an early stage of infection. These findings support a model in which extracellular 2’-3’cGAMP uptake prior to infection functions as an antiviral mechanism in THP-1 cells, as well as the more biologically relevant SH-SY5Y neuroblastoma cell line by inhibiting viral replication at an immediate early gene level.

**Figure 2.**
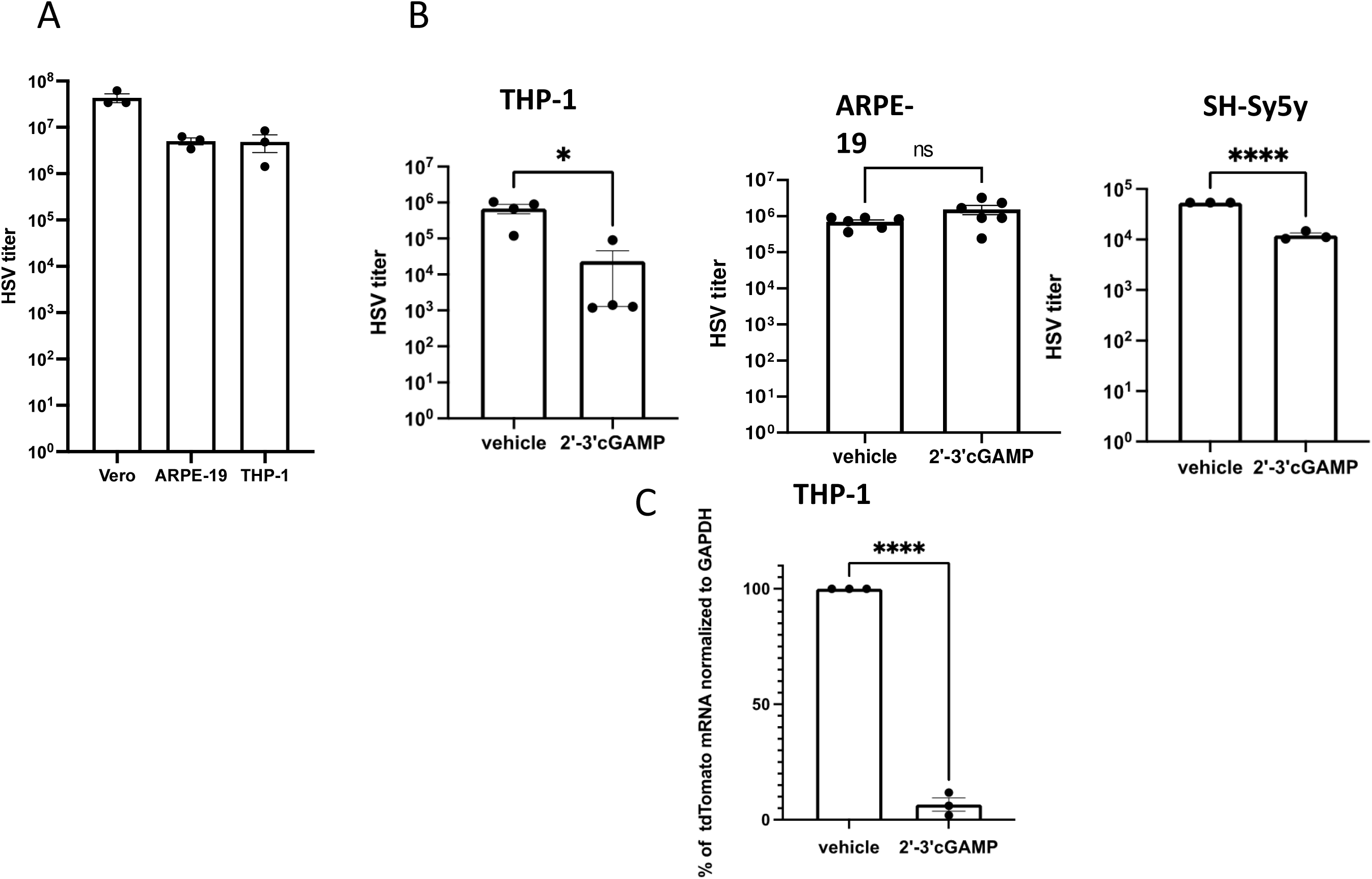
Exogenous 2’-3’cGAMP inhibits HSV-1 replication in THP-1 and SH-SY5Y cells. (A) Vero , ARPE-19 and THP-1 cells were infected with HSV-1 US11eGFP Patton strain (moi=0.5) for either 24h (Vero) or 48h (ARPE-19 and THP-1) after which supernatant was collected and TCid50 assay performed to quantify viral replication. Cells were collected after an additional 24 hours for analysis of viral replication in Vero cells. (B) TCID50 assays were performed in Vero cells on supernatants collected from HSV-1 infected THP-1, ARPE-19 and SH-SY5Y neuroblastoma cells (moi=0.5) with or without 2’-3’cGAMP pre-treatment. (C) THP-1 cells were pre-treated with exogenous 2’-3’cGAMP for 24 h then infected with tdTomato HSV-1 (F-strain) (moi =0.5) and RT-QPCR was performed for both tdTomato and the housekeeping gene GAPDH. n=3. ****= pValue <0.0001, ***=0.001, **=0.01, *=0.05

### Uptake of exogenous 2’-3’cGAMP requires SLC19A1 to establish an antiviral state

Our data reveal extracellular 2’-3’cGAMP treatment of THP-1 cells leads to the activation of an antiviral pathway (Fig 2). Given these findings, we next asked whether SLC19A1-mediated import of 2’-3’cGAMP is required to induce this antiviral response. To test the impact of SLC19A1 transport on viral transcription, we utilized the anti-inflammatory drug, sulfasalazine, which inhibits the uptake of CDNs including 2’-3’cGAMP (31). We first validated that treatment with the drug blocked 2’-3’cGAMP treatment mediated IFNB activation (Supplementary Figure 3) using concentrations we confirmed were not cytotoxic to the cells (Fig 3B). We then pre-treated THP-1 cells with the inhibitor for 4h, after which we added 2’-3’cGAMP to the cultures for 24h. We then infected the cells with HSV-1 US11eGFP Patton, and quantified the expression of the viral IE gene, *ICP27* at 24hpi. In support of our above findings, treatment with 2’-3’cGAMP prior to infection resulted in a decrease of *ICP27* mRNA levels (Fig 3C). However, sulfasalazine pre-treatment alone prior to infection did not significantly impact *ICP27* transcript levels compared to vehicle-treated cells (Fig 3C). Importantly, the cultures treated with both sulfasalazine and 2’-3’cGAMP prior to infection displayed *ICP27* transcript levels similar to that of vehicle treated infected cells (Fig 3C). These data suggest that SLC19A1 inhibition leads to increased viral IE gene expression, indicating the inhibition of this antiporter functions to limit antiviral signaling upon exogenous 2’-3’cGAMP stimulation.

**Figure 3.**
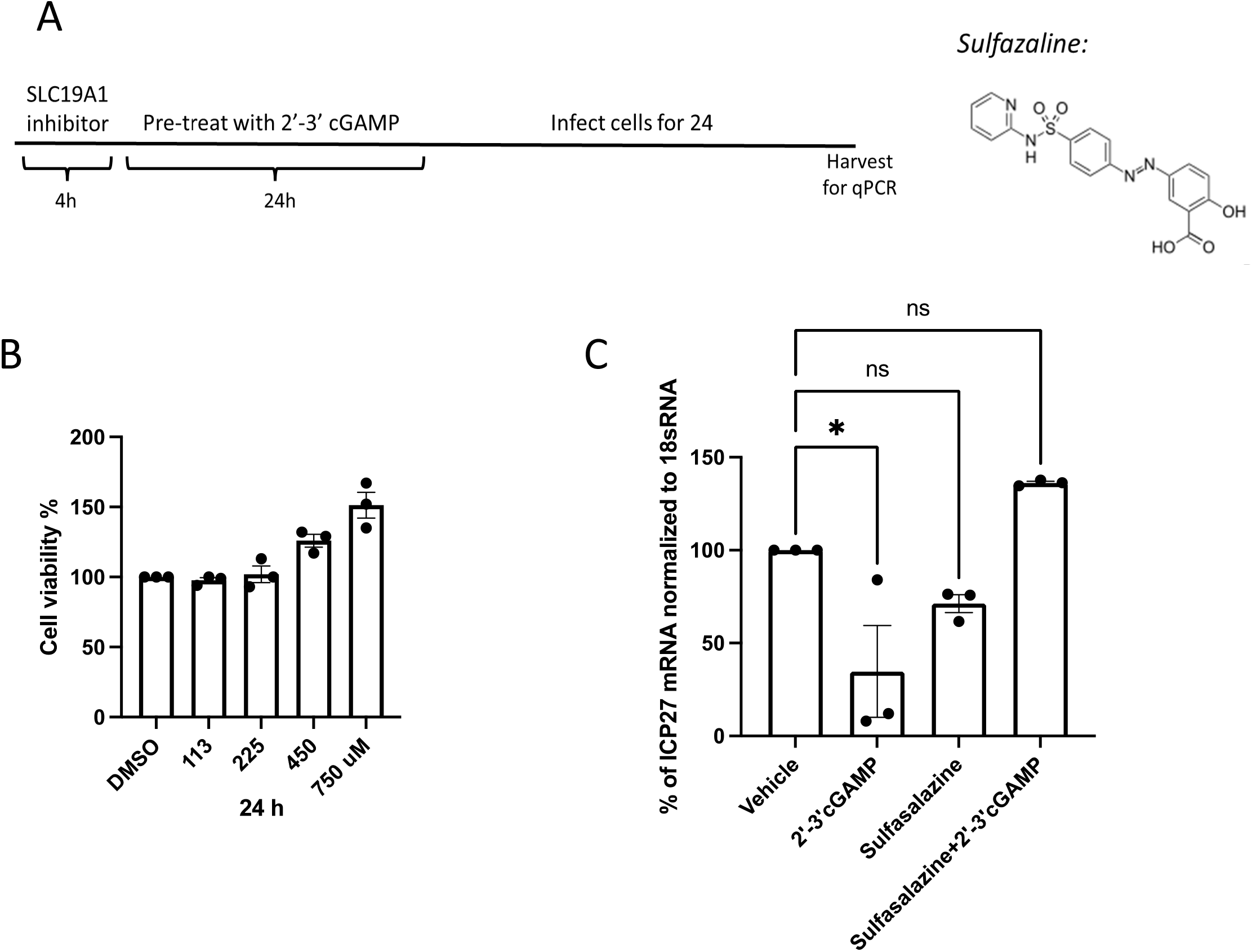
Inhibition of SLC19A1 rescues the antiviral state established by exogenous 2’-3’cGAMP treatment. (A) A timeframe of the treatment conditions is shown. THP-1 cells were pre-treated with SLC19A1 inhibitor, Sulfasalazine (700uM), for 4 hours with or without 2’-3’cGAMP (20ug/ml) prior to infection with HSV-1 UL11eGFP Patton strain (moi=0.3). (B) A cytotoxicity assay was performed using an MTT assay 24h. (C) ICP27 mRNA levels were measured by RT-qPCR. Fold-change levels are normalized to the 18s rRNA house-keeping gene mRNA. n=3. ****= pValue <0.0001, ***=0.001, **=0.01, *=0.05

As drug treatments have the potential to contribute to off-target effects, we used a more focused genetic approach to fully define the requirement for SLC19A1 in inducing an antiviral response. To this end, we utilized CRISPR engineered SLC19A1 KO THP-1 cells (a kind gift from Dr. Rutger Luteijn, Utrecht University) (14), which lack detectable SLC19A1 expression (Fig 4A). We then pre-treated the parental or KO THP-1 cells with exogenous 2’-3’cGAMP for 24 h, at which point we infected each cell line with HSV-1 for an additional 24 h. We then quantified the production of infectious virus from these conditions. Cultures treated with exogenous 2’-3’cGAMP displayed a reduction in HSV-1 production in parental THP-1 cells when compared to untreated counterpart cultures (Fig 4B), similar to our earlier observations (Fig 2B). However, there was no significant difference observed in viral growth upon 2’-3’cGAMP treatment within the SLC19A1 KO THP-1 cells when compared to vehicle treatment (Fig 4B). This is in line with previous reports that 2’-3’cGAMP uptake is impeded in the KO cells (14). Coupled with our data in Fig 3, these results underscore the role of SLC19A1 in promoting an antiviral state in the presence of exogenous 2’-3’cGAMP.

**Figure 4.**
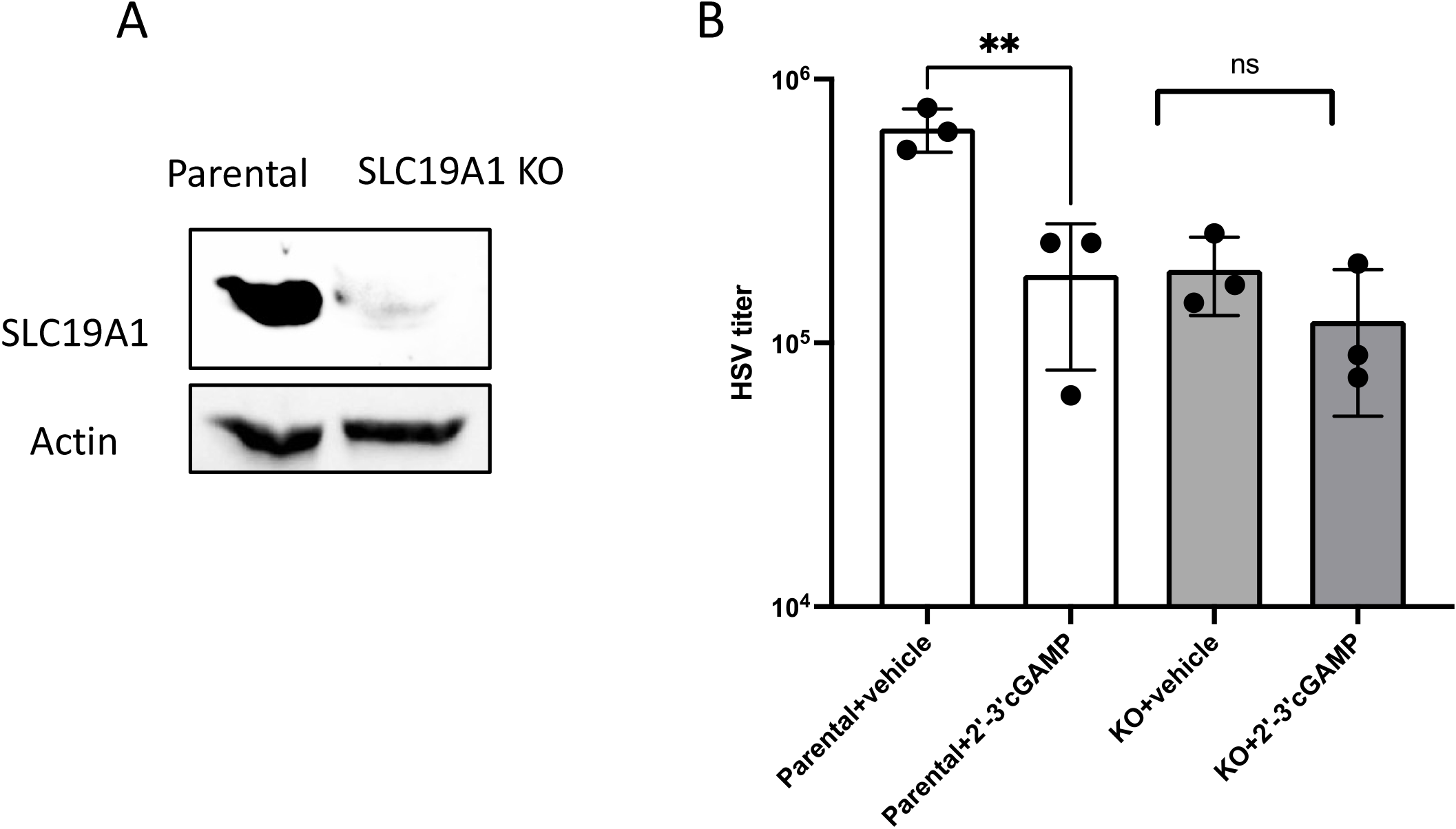
Genetic knockout of SLC19A1 restores HSV1 replication upon treatment with exogenous 2’-3’cGAMP. (A) SLC19A1 expression within parental and KO THP-1 cells were monitored by Western blot using antibodies specific for SLC19A1 and Actin. (B) Parental THP-1 and SLC19A1 KO THP-1 cells were pre-treated with 2’-3’cGAMP (20ug/ml) for 24 hrs followed by infection with HSV-1 (moi=0.3) for an additional 24h. Viral replication was monitored by TCID50 assay using Vero cells. n=3 ****= pValue <0.0001, ***=0.001, **=0.01, *=0.05

### SLC19A1 transcript and protein levels are decreased in HSV-1 infected cells

Given the important role in SLC19A1 establishing an antiviral state, we reasoned HSV-1 likely mounts a defense to counter the extracellular 2’-3’cGAMP-induced viral inhibition. We hypothesized that similar to the virus inhibiting multiple stages of STING activation, HSV-1 may directly target the levels of SLC19A1 to reduce its ability to uptake 2’-3’cGAMP from the extracellular space, thereby blocking a robust immune response. To this end, we evaluated SLC19A1 gene and protein expression in infected cells over time. Upon HSV-1 infection, we observed a significant decrease of *SLC19A1* RNA transcripts over time in both THP-1 and ARPE-19 infected cells when compared to mock-infected cells, beginning as early as 8 hours post-infection (hpi) in THP-1 (Fig 5A) and ARPE-19 cells (Fig 5B), with peak inhibition occurring at 24 hours. Consistent with the transcript data, we observed a significant decrease in total SLC19A1 protein levels in THP-1 cells by 48 hpi, which was not further decreased upon increasing the multiplicity of infection (MOI) (Fig 5C). In contrast, infected ARPE-19 cells did not display reduced SLC19A1 protein levels over the 48-hour time course (Fig 5D). This difference may be explained by differential levels of antiporter expressed on the surface of each cell type or may be due to cell type specific effects of HSV-1 virion host shutoff (VHS) protein, which inhibits host translation. We thus wanted to test if reduction of the total SLC19A1 protein levels we observed during infection are also altered in the cell surface expression where the transporter functions. To this end, THP-1 or ARPE-19 cells were infected with HSV-1 UL11eGFP Patton, followed by monitoring SLC19A1 surface expression by flow cytometry over a time-course of infection ranging from 4h to 48h. We found that HSV-1 infection of THP-1 cells resulted in a significant decrease of SLC19A1-specific cell surface expression as quantified by mean fluorescent intensity values (MFI), thus revealing a downregulation in SLC19A1 surface levels as early as 4 h (Fig 6A, B), a time frame earlier than that observed in our western blot analysis of total protein (Fig 5). Further, we observed a similar reduction in SLC19A1 surface staining in infected ARPE-19 cells, although the kinetics of this decrease was delayed (Fig 6C, D). Importantly, there was a comparably lower amount of SLC19A1 expression on the surface of uninfected ARPE-19 cells compared to THP-1 cells, which correlates with our data above showing a reduced induction of *IFNB* and robust viral infection in ARPE-19 cells despite extracellular 2’-3’cGAMP treatment (Fig 1B and 2B). Furthermore, our results reveal that this effect is specific for infected cells, as gating for the uninfected population showed no change in the SLC19A1 cell surface expression (Supplementary Figure 4). Overall, these data show that HSV-1 infection reduces SLC19A1 RNA, protein, and cell surface expression in THP-1 cells, suggesting a novel mechanism by which HSV-1 undermines the STING pathway.

**Figure 5.**
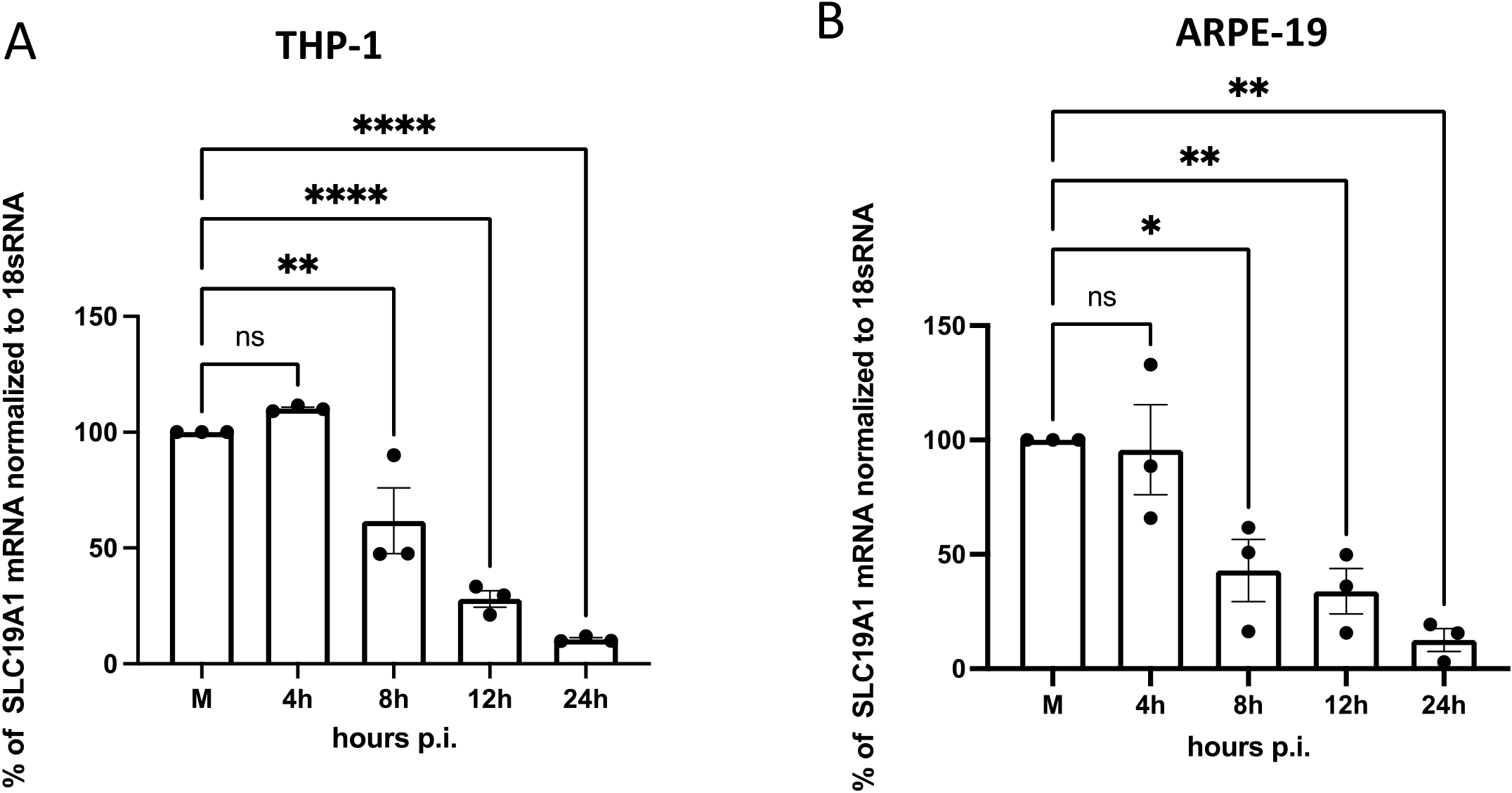
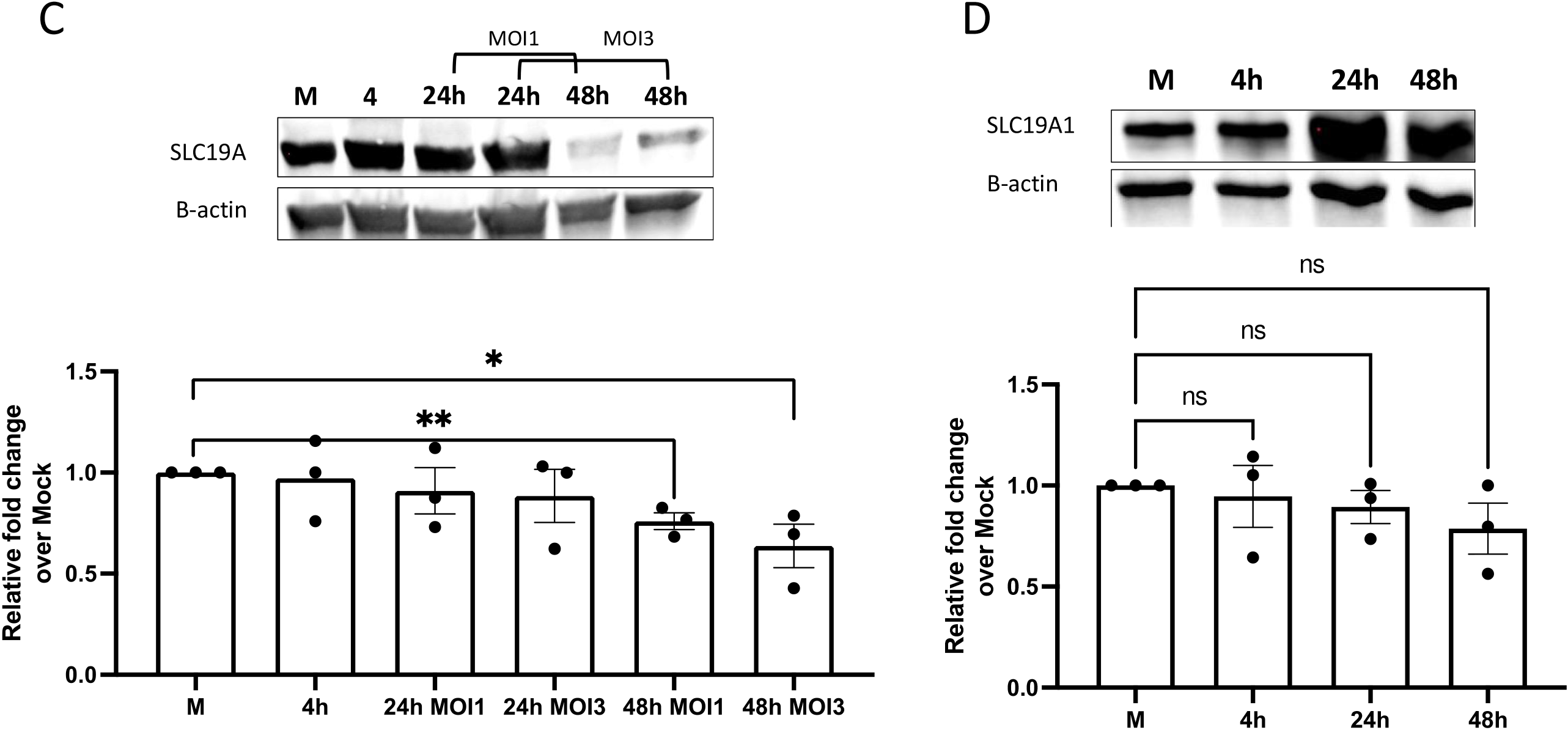
SLC19A1 expression is significantly decreased in THP-1 and to a lesser degree, in ARPE-19 cells infected by HSV-1. (A & B) THP-1 and APRE-19 cells were infected and total RNA was isolated at the indicated time-course (moi=0.5). RT-qPCR was performed, and relative fold change levels were normalized to 18s rRNA house-keeping gene . (C & D) THP-1 and APRE-19 cells were infected and protein lysates were isolated at the indicated times (moi=0.5). Western blot analyzes was performed and fold change was calculated after normalization to ß-actin over mock uninfected cells. Corresponding densitometry analyses is shown in panel C and D. ****= pValue <0.0001, ***=0.001, **=0.01, *=0.05

**Figure 6.**
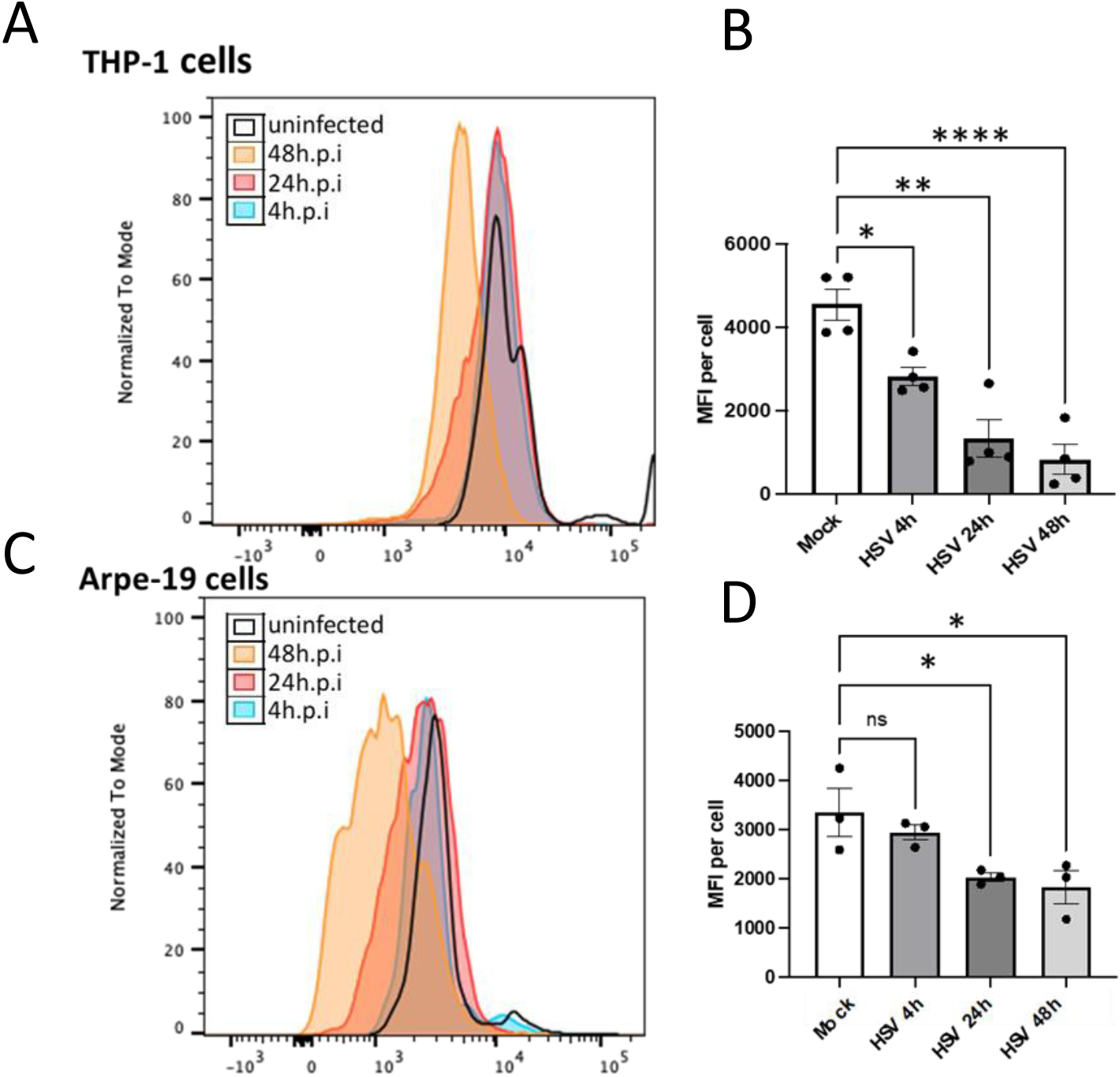
SLC19A1 cell surface expression is significantly decreased in THP-1 and ARPE-19 cells upon infection by HSV-1. (A) THP-1 or (B) ARPE-19 cells were infected with HSV-1 US11eGFP Patton strain (moi=0.5) for the indicated time course. SLC19A1 cell surface expression was assessed by flow cytometry and the cells were gated for living and GFP+ (infected) cells only. Corresponding MFI values are quantified. n=3 . ****= pValue<0.0001, **=0.01, *=0.05

### Viral immediate early and/or early gene transcription is required for the observed decrease of SLC19A1 expression

We next evaluated the stage of the HSV-1 life cycle that contributes to the decrease in SLC19A1. To this end, we infected THP-1 cells with HSV-1 or UV-inactivated HSV-1. UV inactivation induces thymidine dimers in the viral DNA, preventing *de novo* viral transcription (32,33). In parallel, we also treated HSV-1-infected cultures with phosphonoacetic acid (PAA), which blocks the viral polymerase, thus in turn inhibiting viral DNA replication (34), which is necessary for viral late gene expression (Supplementary Figure 5). Importantly, these treatments do not inhibit the delivery or functions of the viral tegument proteins. Upon infection, we observed that infected cultures treated with PAA showed a significant decrease in SLC19A1 protein levels, similar to what was observed in cells infected with WT virus (Fig 7), suggesting that the reduction of the host protein was not dependent on the expression of a late viral gene product. Importantly, there was no significant reduction in SLC19A1 levels when comparing mock-infected and UV-inactivated virus infected cells (Fig. 7), which suggests that the downregulation of SLC19A1 requires a step in the viral replication cycle prior to late gene expression, but after virion entry. Additionally, as SLC19A1 expression is not attenuated with UV-inactivated virus, the repression is not mediated by HSV-1 tegument delivered proteins. Thus, the downregulation of SLC19A1 in HSV-1 infected cells is mediated by an immediate-early and/or early viral gene.

**Figure 7.**
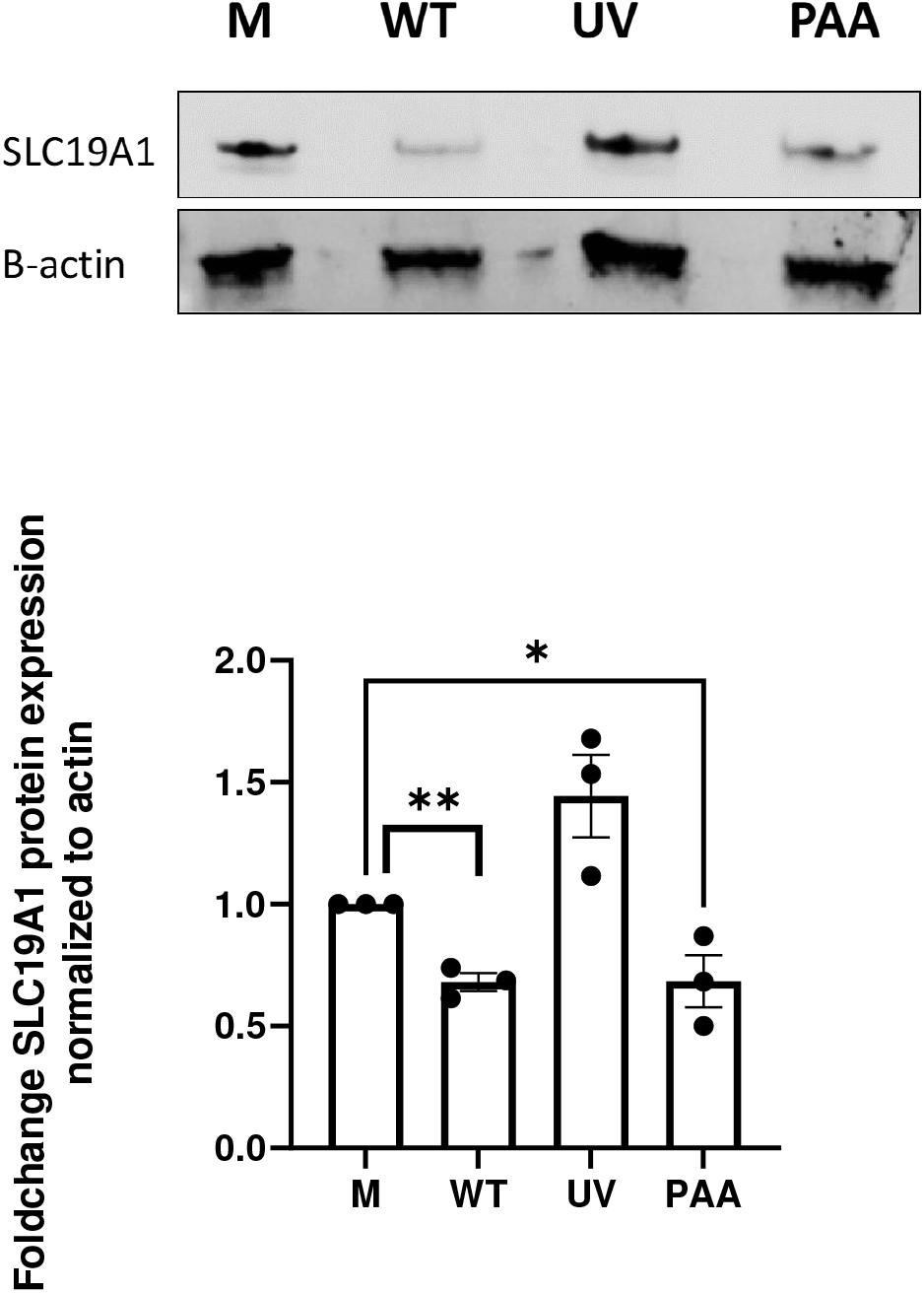
HSV-1 immediate early and/or early gene expression is required to mediate SLC19A1 downregulation in infected cells. THP-1 cells were infected with UV inactivated HSV-1 or treated with PAA (300ug/ml) along with infection with HSV-1(moi=0.5) for 48h, then Western blot analysis was performed with an SLC19A1 specific primary antibody. Densitometry analysis of three Western blots and a representative blot is shown. Fold change levels are normalized to ß-actin and uninfected cells. ****= pValue <0.0001, ***=0.001, **=0.01, *=0.05

## DISCUSSION

We examined extracellular 2’-3’cGAMP as a potential messenger that can induce an antiviral response thereby limiting HSV-1 replication. We discovered that in the THP-1 cell line, widely used for investigating the STING pathway,(35–37) adding the cyclic di-nucleotide, 2’-3’cGAMP, to the medium led to interferon production and reduced viral replication (Figs 1 & 2). These results indicate that not only *de novo* cGAS synthesized 2’-3’cGAMP, but also extracellular 2’-3’cGAMP, initiates an antiviral state in a dose-dependent manner, which aligns with our hypothesis based on findings in tumor cells (13,14). Our data is supported by a recent publication showing increased IFN levels in THP-1 cells when treated with 2’-3’cGAMP (38,39).

By employing an SLC19A1 inhibitor, sulfasalazine, and genetic knockout of the transporter, we demonstrated SLC19A1 is required for the exogenous 2’-3’cGAMP-mediated inhibition of viral gene expression. Abrogating SLC19A1 resulted in higher viral replication with the addition of 2’-3’cGAMP compared to parental cells treated solely with 2’-3’cGAMP, suggesting SLC19A1 operates to support an antiviral state, necessitating the virus to counter this mechanism (Figs 3 and 4). Further, these findings reveal HSV-1 counters this antiviral response by limiting SLC19A1 RNA, total protein, and cell surface expression, thereby favoring successful viral replication. It is possible other newly identified CDN transporters (39) could also play a role in HSV-1 replication. Whether this is the case, and whether HSV-1 also targets these other transports as an added countermeasure, warrants further investigation.

Previous reports indicated that HSV-1 targets various components of the STING pathway to sustain its replication (40–43). Consistent with these findings, we found a novel means by which HSV-1 subverts STING signaling by attenuating SLC19A1 expression, ultimately reducing its expression and availability on the cell surface (Figs 5 & 6). Importantly, the kinetics of reduced SLC19A1 cell surface expression in infected THP-1 cells (∼4hpi) suggests that the virus rapidly reduces this protein thereby undermining a non-canonical activation of the STING pathway. Our findings also suggest either an HSV-1 IE and/or early gene is responsible for SLC19A1 protein suppression in THP-1 cells (Fig 7). Unraveling the specific genes responsible for interfering with STING signaling via this novel mechanism could enhance the understanding of this novel mechanism.

For successful replication, viruses must counteract as many of the host’s antiviral responses as possible. Herpesviruses such as EBV, KSHV, HCMV, HSV-1, and HSV-2 have all developed unique strategies to subvert one of the host’s most central immune defense mechanisms in response to cytosolic DNA, the STING pathway. They achieve this feat at both the sensing and signaling levels. Several viral proteins, including HSV-1-encoded ICP0 and ICP4, KSHV-encoded vIRF1, ORF52, and LANA, EBV ORF52, and HCMV-encoded pp71, pp65 and UL36a, target either the STING protein itself, cGAS, or the signaling process through the STING-TBK1 axis (44–47,47–50). Our findings now add an additional STING target inhibited by the virus.

The significance of the STING pathway extends beyond virally infected cells to other cellular processes. For instance, cancer cells also harbor cytoplasmic DNA as a result of genomic instability (51). Cancer cells, much like virally infected cells, display high levels of STING activation and promote an anti-tumor state by producing interferons (51–53). Upon detecting cytosolic DNA, cGAS generates *de novo* 2’-3’cGAMP that activates STING (54) . While the role of 2’-3’cGAMP as an intracellular messenger is well-established (16,18), recent studies have highlighted its emerging role as an “immunotransmitter” in the tumor microenvironment. Cancer cells release 2’-3’cGAMP into the extracellular space, allowing communication to neighboring cells that induce a signaling event, thus propagating the response in a paracrine manner (13,15). Thus, STING agonists, including 2’-3’cGAMP, are being explored as promising adjuvant therapies due to their ability to enhance the host’s immune response to cancer cells (55–57). Our study adds to the potential of exploiting SLC19A1 import of exogenous 2’-3’cGAMP to limit viral infections.

In conclusion, we identified a novel mechanism using the transporter, SLC19A1, by which HSV-1 counters the antiviral STING pathway. Much like what is observed in the interferon responses of infected cells (58), we hypothesize 2’-3’cGAMP is released from infected cells and influences neighboring cells in a paracrine manner (Fig 8). In turn, HSV-1 inhibits this mechanism by reducing the SLC19A1 transporter, thereby inhibiting its ability to internalize extracellular 2’-3’cGAMP. Future investigations are aimed at understanding the detailed biological mechanisms underlying this antiviral countermeasure, characterizing the HSV-1 encoded factors that mediate this inhibition and investigating how ubiquitous this response is among pathogens.

**Figure 8.**
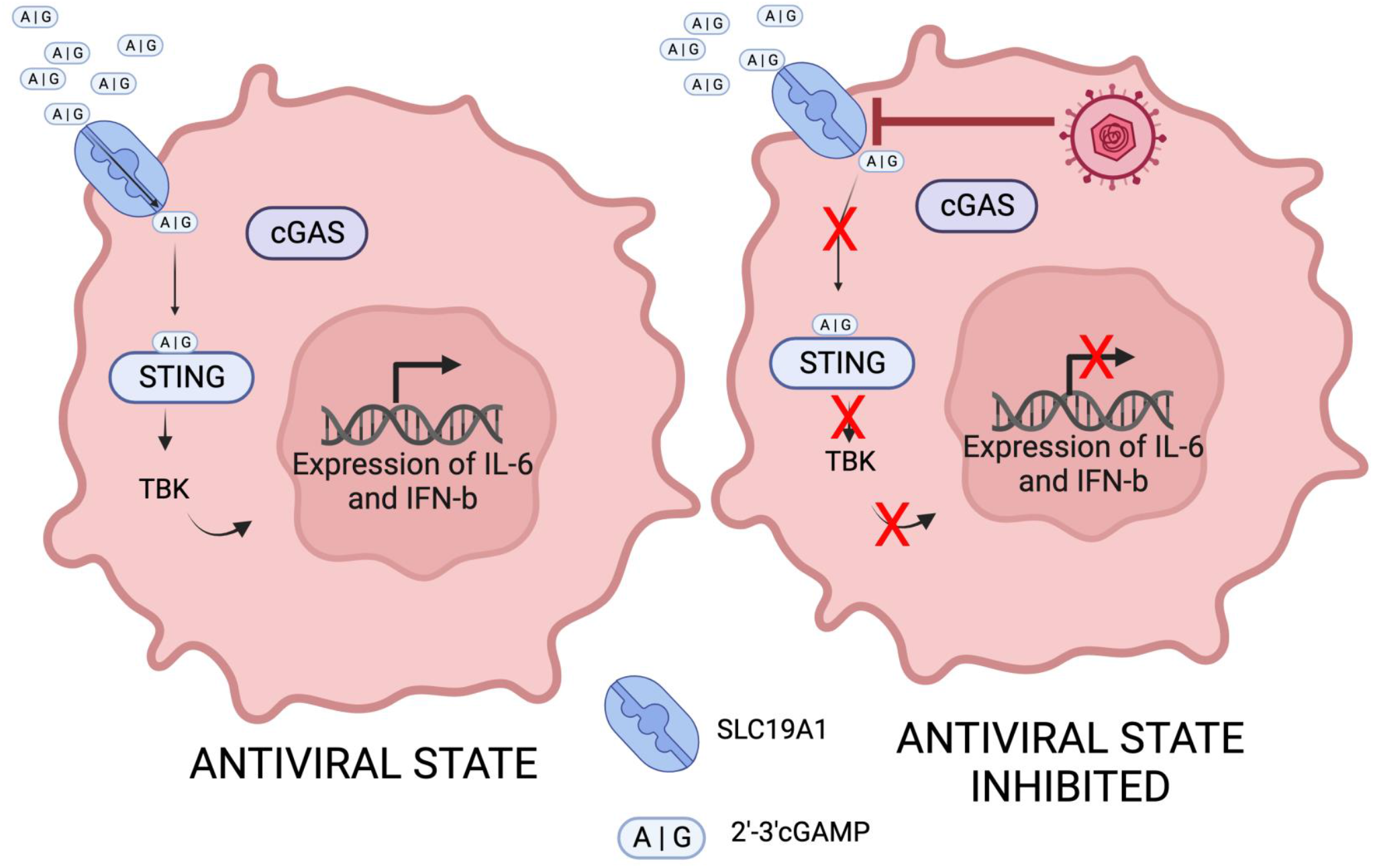
Model of HSV-1 downregulation of SLC19A1 undermines the antiviral state established after the addition of extracellular 2’-3’cGAMP. (Left) Exogenous 2’-3’cGAMP import by SLC19A1 induces STING via a cGAS independent mechanism thereby inducing the expression of host cytokines resulting in an antiviral state. **(Right)** HSV-1 infection reduces the cell surface expression of SLC19A1 thereby limiting the import of 2’-3’cGAMP resulting in a diminished antiviral response.

## Materials and Methods

### Cells and culture conditions

THP-1 monocytic cells (ATCC) were cultured in RPMI 1640 (10-500p, Clevelend Clinic Media Core, Cleveland Clinic, OH) with 10% fetal bovine serum (FBS), 1% glutamine, and 1% Pen-Strep, 10 μg/ml gentamicin, 0.18 uM beta-mercaptoethanol, and 20 mM HEPES. Cells were maintained at a density of 0.5 million cells per ml. ARPE-19 cells (ATCC) (passage 34 – 40) were cultured in DMEM: F12 medium (1:1 ratio) (11-500p, Clevelend Clinic Media Core), supplemented with 10% FBS, 1% glutamine, and 1% Pen-Strep and were maintained under 90% confluency. Vero cells (ATCC) were grown in 1x DMEM (11-500p, Clevelend Clinic Media Core) supplemented with 10% fetal bovine serum (FBS), 1% glutamine, and 1% Pen-Strep and were maintained under 90% confluency. SLC19A1 KO THP-1 cells with the corresponding parental cell lines were generously provided by Dr. Rutger Luteijn (Utrecht,Amsterdam, NT) and cultured under the same conditions as described above for THP-1 cells. Regular microscopic examinations and mycoplasma testing (Myco Strip, Invivogen) ensured cells were free of contamination, and morphological changes indicative of differentiation or cellular stress were monitored. All cells were maintained at 37°C in an atmosphere containing 5% CO_2_.

The SHSY5Y neuroblastoma cell line was grown and differentiated by the Shipley protocol for 18 days (29). Cells were grown in Eagle’s Minimum Essential Medium that contained 15%FBS, 1%P/S, 2 mM Glutamine, (99BJ500CUSTp,Clevelend Clinic) and differentiated using 2.5% FBS neurobasal media (1X B27, 20 mM KCl, 1% Pen-Strep, 2 mM Glutamax (Thermo Fisher) 50 ng/mL Brain-derived neurotrophic factor (BDNF), 2 mM dibutyryl cyclic AMP (db-cAMP), and 10 μM retinoic acid) for 7 days (d). After 7d, the media was removed, and cells were plated in laminin pre-coated dishes in EMEM media containing 1% FBS. Finally, cells were cultured for an additional 11 d in neurobasal medium (Fisher Scientific) and 10 uM retinoic acid, with media changes every 3d. Differentiated cells were split at 70% confluency and passaged to a maximum of 10-15 passages.

### Viruses and Viral Infection

Viral stocks were grown in Vero cells (ATCC) until complete cytopathic effect was observed. Supernatants were harvested, aliquoted, and stored at -80°C to maintain virus viability. Viral stocks were titered by TCID_50_ assay on Vero cells. The BAC HSV-1 Strain F expressing tdTomato driven by the SV40 promoter was generously provided by Dr. Greg Smith (30). The HSV-1 eGFP US11eGFP Patton Strain was a kind gift from Dr. Ian Mohr (28). Where indicated, virus was UV-inactivated using a UV STRATAlinker 2400 box (Stratagene, San, Diego, CA). To block viral late gene expression, viral polymerase was inhibited using PAA (300 μg) (Millipore Sigma).

### TCID50 Assay

Virus stock titers are conveyed as TCID50. To quantify infectious viral particles, we utilized the protocol of the Reed laboratory (59). Vero cells (90% confluency) were plated on 96-well plates and infected with serial dilutions of the virus. After 48h wells demonstrating fluorescence were counted, and the end-point dilution was determined to calculate the TCID50 titer (PFU/ml).

### Cell Viability

Cell viability post-drug treatment was measured via both MTT or WST assays according to manufactorer’s protocol (Promega) and automatic cell counter measurements (Luna II automatic cell counter), using Trypan blue for live-dead discrimination, providing an additional metric for assessing cell health post-drug treatment for both 2’-3’cGAMP (Invivogen) and Sulfasalazine (Cayman Chemical).

### Protein Analyses

Protein was harvested using lysis buffer (Pierce RIPA, Thermo Scientific) complemented with complete Protease and Phosphatase inhibitor (Pierce, Thermo Scientific). Equal protein quantities were separated on a 10% SDS-PAGE gel, transferred to nitrocellulose via semi-dry transfer, and after blocking with 5% BSA or Everything Blocking Buffer (Biorad) for an hour at room temperature, probed with antibodies to SLC19A1 (Santa Cruz. D-4 IgG1clone, 1:800 dilution in blocking buffer) or β-actin rhodamine (Biorad,1:5000 in blocking buffer) at 4°C overnight. Following incubation with a secondary mouse antibody (Cell signaling,1:1000 dilution in blocking buffer) at room temperature, membranes were washed in buffer containing 0.1% Tween-20 three times. Membranes were visualized using ECL substrate on a ChemiDoc MP (Biorad). To ensure equal loading between samples, before loading on SDS-PAGE, protein concentrations were determined using the Bradford Protein Assay (Thermo Scientific).

### Quantification of RNA

RNA quantifications were analyzed via Real Time - quantitative PCR (RT-qPCR) in triplicate, employing the Power SYBR Green Master Mix (BioRad) or Taqman assay method on the Bio-Rad CFX Connect real-time PCR device. Fluorescent primers specific to SLC19A1 (Hs00953344_m1), IFNB (Hs01077958_s1), GAPDH (4326317E), 18sRNA (Hs03928990 _g1), were acquired from Thermo Fisher. EGFP and ICP27 primers for SYBR Green and fluorescent applications were purchased from IDT.

eGFP Forward Primer: 5’-GATCGACTACGCGACCCTTG-3’

eGFP reverse Primer: 5’-GCAGACACGACTCGAACACT-3’

ICP27 Forward Primer: 5’-ACCACTACCTGAGCACCCAGTC-3’

ICP27 Reverse Primer: 5’-GTCCATGCCGAGAGTGATCC-3’

SLC19A1 Forward Primer: 5’-GGACAGGATCAGGAAGTACACG-3’

SLC19A1 Reverse Primer: 5’-CTTTGCCACCATCGTCAAGACC-3’

RNA was harvested using TRIzol Reagent (Invitrogen), ensuring RNase-free conditions, following the manufacturer’s guidelines. Following isopropanol precipitation, equal quantities of RNA (1 μg) were either reverse transcribed to cDNA using the TaqMan kit (Applied Biosystems) or directly used with the ONE STEP TAQ QPCR kit (Promega). For RT-qPCR assays, either the Power SYBR Green Master Mix (Applied Biosystems) or fluorescent probes were used to quantify gene expression using a CFX Connect QPCR machine (Biorad). GAPDH or 18sRNA served as a host reference gene.

### Flow Cytometry

After HSV-1 US11eGFP infection, ARPE-19 or THP-1 cells were stained using the LiveDead Zombie (BioLegend) and SLC19A1 PE antibody (D-4 clone, Santa Cruz) for 15 minutes at room temperature. Cells were then washed with FACS buffer, then resuspended in FACS buffer supplemented with 5% FBS and analyzed on a LSR Fortessa Flow Cytometer for cell surface staining. Cells were gated for size/shape, viability, and infection (by GFP) and surface expression of SLC19A1 was determined relative to controls using FlowJo software. Fluorescent minus one controls were integrated into the analysis. To correct for spectral overlap between fluorochromes, compensation was performed using single-stained controls, ensuring accurate interpretation of multi-color flow cytometry data.

### Statistics

All results represent a minimum of three independent biological replicates. Data were processed using t-tests, one-way or two-way ANOVA, and presented as means ± standard errors. Results were plotted using GraphPad Prism software.

## ACKNOWLEDGMENTS

We are grateful to Dr. Rutger Luteijn for his kind help and the SLC19A1 KO cells he provided for our research. Also, I would like to thank Victoria R Zoccoli for her help with performing Flow-analyses and our lab members Justin Cox and Megan Carter for their support and advice for the project.

E.M. acknowledges support from the National Institutes of Health (Award Nos. 5R01AI155979 and 5R01AG076007).

Z.S. and E.M. conceived of the study. Z.S. performed all the experiments. Z.S. wrote the first manuscript draft. Z.S. and E.M. reviewed and edited the manuscript.

## REFERENCES

1. Ishikawa H, Barber GN. STING is an endoplasmic reticulum adaptor that facilitates innate immune signalling. Nature. 2008 Oct 2;455(7213):674–8.

2. Xiao TS, Fitzgerald KA. The cGAS-STING pathway for DNA sensing. Mol Cell. 2013 Jul 25;51(2):135–9.

3. Zhong B, Yang Y, Li S, Wang YY, Li Y, Diao F, et al. The adaptor protein MITA links virus-sensing receptors to IRF3 transcription factor activation. Immunity. 2008 Oct 17;29(4):538–50.

4. Diamond MS. IFIT1: A dual sensor and effector molecule that detects non-2′-O methylated viral RNA and inhibits its translation. Cytokine Growth Factor Rev. 2014 Oct;25(5):543–50.

5. Schoggins JW, MacDuff DA, Imanaka N, Gainey MD, Shrestha B, Eitson JL, et al. Pan-viral specificity of IFN-induced genes reveals new roles for cGAS in innate immunity. Nature. 2014 Jan 30;505(7485):691–5.

6. Ablasser A, Goldeck M, Cavlar T, Deimling T, Witte G, Röhl I, et al. cGAS produces a 2’-5’-linked cyclic dinucleotide second messenger that activates STING. Nature. 2013 Jun 20;498(7454):380–4.

7. Diner EJ, Burdette DL, Wilson SC, Monroe KM, Kellenberger CA, Hyodo M, et al. The innate immune DNA sensor cGAS produces a noncanonical cyclic dinucleotide that activates human STING. Cell Rep. 2013 May 30;3(5):1355–61.

8. Yin Q, Tian Y, Kabaleeswaran V, Jiang X, Tu D, Eck MJ, et al. Cyclic di-GMP sensing via the innate immune signaling protein STING. Mol Cell. 2012 Jun 29;46(6):735–45.

9. Verpooten D, Ma Y, Hou S, Yan Z, He B. Control of TANK-binding Kinase 1-mediated Signaling by the γ134.5 Protein of Herpes Simplex Virus 1. J Biol Chem. 2009 Jan 9;284(2):1097–105.

10. Boutell C, Sadis S, Everett RD. Herpes Simplex Virus Type 1 Immediate-Early Protein ICP0 and Its Isolated RING Finger Domain Act as Ubiquitin E3 Ligases In Vitro. J Virol. 2002 Jan;76(2):841–50.

11. Christensen MH, Jensen SB, Miettinen JJ, Luecke S, Prabakaran T, Reinert LS, et al. HSV-1 ICP27 targets the TBK1-activated STING signalsome to inhibit virus-induced type I IFN expression. The EMBO Journal. 2016 Jul;35(13):1385–99.

12. Sun L, Wu J, Du F, Chen X, Chen ZJ. Cyclic GMP-AMP synthase is a cytosolic DNA sensor that activates the type I interferon pathway. Science. 2013 Feb 15;339(6121):786–91.

13. Carozza JA, Böhnert V, Nguyen KC, Skariah G, Shaw KE, Brown JA, et al. Extracellular cGAMP is a cancer cell-produced immunotransmitter involved in radiation-induced anti-cancer immunity. Nat Cancer. 2020 Feb;1(2):184–96.

14. Luteijn RD, Zaver SA, Gowen BG, Wyman SK, Garelis NE, Onia L, et al. SLC19A1 transports immunoreactive cyclic dinucleotides. Nature. 2019 Sep;573(7774):434–8.

15. Carozza JA, Cordova AF, Brown JA, AlSaif Y, Böhnert V, Cao X, et al. ENPP1’s regulation of extracellular cGAMP is a ubiquitous mechanism of attenuating STING signaling. Proceedings of the National Academy of Sciences. 2022 May 24;119(21):e2119189119.

16. Li T, Cheng H, Yuan H, Xu Q, Shu C, Zhang Y, et al. Antitumor Activity of cGAMP via Stimulation of cGAS-cGAMP-STING-IRF3 Mediated Innate Immune Response. Sci Rep. 2016 Jan 12;6(1):19049.

17. Carozza JA, Brown JA, Böhnert V, Fernandez D, AlSaif Y, Mardjuki RE, et al. Structure-aided development of small molecule inhibitors of ENPP1, the extracellular phosphodiesterase of the immunotransmitter cGAMP. Cell Chem Biol. 2020 Nov 19;27(11):1347–1358.e5.

18. Ablasser A, Schmid-Burgk JL, Hemmerling I, Horvath GL, Schmidt T, Latz E, et al. Cell intrinsic immunity spreads to bystander cells via the intercellular transfer of cGAMP. Nature. 2013 Nov 28;503(7477):530–4.

19. Bridgeman A, Maelfait J, Davenne T, Partridge T, Peng Y, Mayer A, et al. Viruses transfer the antiviral second messenger cGAMP between cells. Science. 2015 Sep 11;349(6253):1228–32.

20. Ritchie C, Cordova AF, Hess GT, Bassik MC, Li L. SLC19A1 Is an Importer of the Immunotransmitter cGAMP. Mol Cell. 2019 Jul 25;75(2):372–381.e5.

21. Matherly LH, Goldman DI. Membrane transport of folates. Vitam Horm. 2003;66:403–56.

22. Zheng C. Evasion of Cytosolic DNA-Stimulated Innate Immune Responses by Herpes Simplex Virus 1. J Virol. 2018 Mar 15;92(6):e00099–17.

23. Zhu H, Zheng C. The Race between Host Antiviral Innate Immunity and the Immune Evasion Strategies of Herpes Simplex Virus 1. Microbiol Mol Biol Rev. 2020 Nov 18;84(4):e00099–20.

24. Lin Y, Zheng C. A Tug of War: DNA-Sensing Antiviral Innate Immunity and Herpes Simplex Virus Type I Infection. Front Microbiol. 2019;10:2627.

25. Mosca JD, Pitha PM. Transcriptional and posttranscriptional regulation of exogenous human beta interferon gene in simian cells defective in interferon synthesis. Mol Cell Biol. 1986 Jun;6(6):2279– 83.

26. Desmyter J, Melnick JL, Rawls WE. Defectiveness of interferon production and of rubella virus interference in a line of African green monkey kidney cells (Vero). J Virol. 1968 Oct;2(10):955–61.

27. Chamma H, Guha S, Laguette N, Vila IK. Protocol to induce and assess cGAS-STING pathway activation in vitro. STAR Protocols. 2022 Jun 17;3(2):101384.

28. Benboudjema L, Mulvey M, Gao Y, Pimplikar SW, Mohr I. Association of the Herpes Simplex Virus Type 1 Us11 Gene Product with the Cellular Kinesin Light-Chain-Related Protein PAT1 Results in the Redistribution of Both Polypeptides. J Virol. 2003 Sep;77(17):9192–203.

29. Shipley MM, Mangold CA, Szpara ML. Differentiation of the SH-SY5Y Human Neuroblastoma Cell Line. J Vis Exp. 2016 Feb 17;(108):53193.

30. Stults AM, Smith GA. The Herpes Simplex Virus 1 Deamidase Enhances Propagation but Is Dispensable for Retrograde Axonal Transport into the Nervous System. J Virol. 2019 Oct 29;93(22):e01172–19.

31. Jansen G, van der Heijden J, Oerlemans R, Lems WF, Ifergan I, Scheper RJ, et al. Sulfasalazine is a potent inhibitor of the reduced folate carrier: implications for combination therapies with methotrexate in rheumatoid arthritis. Arthritis Rheum. 2004 Jul;50(7):2130–9.

32. Mossman KL, Macgregor PF, Rozmus JJ, Goryachev AB, Edwards AM, Smiley JR. Herpes Simplex Virus Triggers and Then Disarms a Host Antiviral Response. J Virol. 2001 Jan;75(2):750–8.

33. Luftig MA. Viruses and the DNA Damage Response: Activation and Antagonism. Annual Review of Virology. 2014;1(1):605–25.

34. Phosphonoacetic Acid - an overview | ScienceDirect Topics [Internet]. [cited 2024 Jan 8]. Available from: https://www.sciencedirect.com/topics/medicine-and-dentistry/phosphonoacetic-acid

35. Qiao Y, Zhu S, Deng S, Zou SS, Gao B, Zang G, et al. Human Cancer Cells Sense Cytosolic Nucleic Acids Through the RIG-I–MAVS Pathway and cGAS–STING Pathway. Frontiers in Cell and Developmental Biology [Internet]. 2021 [cited 2024 Jan 5];8. Available from: https://www.frontiersin.org/articles/10.3389/fcell.2020.606001

36. Ilg T. Investigations on the molecular mode of action of the novel immunostimulator ZelNate: Activation of the cGAS-STING pathway in mammalian cells. Molecular Immunology. 2017 Oct 1;90:182–9.

37. Decout A, Katz JD, Venkatraman S, Ablasser A. The cGAS–STING pathway as a therapeutic target in inflammatory diseases. Nat Rev Immunol. 2021 Sep;21(9):548–69.

38. Zhang L, Wei X, Wang Z, Liu P, Hou Y, Xu Y, et al. NF-κB activation enhances STING signaling by altering microtubule-mediated STING trafficking. Cell Rep. 2023 Mar 28;42(3):112185.

39. Wei X, Zhang L, Yang Y, Hou Y, Xu Y, Wang Z, et al. LL-37 transports immunoreactive cGAMP to activate STING signaling and enhance interferon-mediated host antiviral immunity. Cell Reports. 2022 May;39(9):110880.

40. Krawczyk E, Kangas C, He B. HSV Replication: Triggering and Repressing STING Functionality. Viruses. 2023 Jan 13;15(1):226.

41. Latif MB, Raja R, Kessler PM, Sen GC. Relative Contributions of the cGAS-STING and TLR3 Signaling Pathways to Attenuation of Herpes Simplex Virus 1 Replication. J Virol. 2020 Feb 28;94(6):e01717–19.

42. Wang C, Sharma N, Veleeparambil M, Kessler PM, Willard B, Sen GC. STING-Mediated Interferon Induction by Herpes Simplex Virus 1 Requires the Protein Tyrosine Kinase Syk. mBio. 2021 Dec 21;12(6):e0322821.

43. Pan S, Liu X, Ma Y, Cao Y, He B. Herpes Simplex Virus 1 γ134.5 Protein Inhibits STING Activation That Restricts Viral Replication. J Virol. 2018 Oct 15;92(20):e01015–18.

44. Kalamvoki M, Du T, Roizman B. Cells infected with herpes simplex virus 1 export to uninfected cells exosomes containing STING, viral mRNAs, and microRNAs. Proc Natl Acad Sci U S A. 2014 Nov 18;111(46):E4991–4996.

45. Wu J jun, Li W, Shao Y, Avey D, Fu B, Gillen J, et al. Inhibition of cGAS DNA Sensing by a Herpesvirus Virion Protein. Cell Host Microbe. 2015 Sep 9;18(3):333–44.

46. Nukui M, Roche KL, Jia J, Fox PL, Murphy EA. Protein S-nitrosylation of Human Cytomegalovirus pp71 inhibits its ability to limit STING antiviral responses [Internet]. bioRxiv; 2020 [cited 2023 Sep 8]. p. 2020.01.08.899757. Available from: https://www.biorxiv.org/content/10.1101/2020.01.08.899757v1

47. Zhang G, Chan B, Samarina N, Abere B, Weidner-Glunde M, Buch A, et al. Cytoplasmic isoforms of Kaposi sarcoma herpesvirus LANA recruit and antagonize the innate immune DNA sensor cGAS. Proc Natl Acad Sci U S A. 2016 Feb 23;113(8):E1034–1043.

48. Ma Z, Jacobs SR, West JA, Stopford C, Zhang Z, Davis Z, et al. Modulation of the cGAS-STING DNA sensing pathway by gammaherpesviruses. Proc Natl Acad Sci U S A. 2015 Aug 4;112(31):E4306–4315.

49. Fu YZ, Su S, Gao YQ, Wang PP, Huang ZF, Hu MM, et al. Human Cytomegalovirus Tegument Protein UL82 Inhibits STING-Mediated Signaling to Evade Antiviral Immunity. Cell Host Microbe. 2017 Feb 8;21(2):231–43.

50. Ren Y, Wang A, Zhang B, Ji W, Zhu XX, Lou J, et al. Human cytomegalovirus UL36 inhibits IRF3-dependent immune signaling to counterbalance its immunoenhancement as apoptotic inhibitor. Sci Adv. 9(40):eadi6586.

51. Talens F, Van Vugt MATM. Inflammatory signaling in genomically instable cancers. Cell Cycle. 2019 Jul 10;18(16):1830–48.

52. Woo SR, Fuertes MB, Corrales L, Spranger S, Furdyna MJ, Leung MYK, et al. STING-dependent cytosolic DNA sensing mediates innate immune recognition of immunogenic tumors. Immunity. 2014 Nov 20;41(5):830–42.

53. Kwon J, Bakhoum SF. The Cytosolic DNA-Sensing cGAS–STING Pathway in Cancer. Cancer Discovery. 2020 Jan 9;10(1):26–39.

54. Hussain B, Xie Y, Jabeen U, Lu D, Yang B, Wu C, et al. Activation of STING Based on Its Structural Features. Frontiers in Immunology [Internet]. 2022 [cited 2023 Dec 15];13. Available from: https://www.frontiersin.org/articles/10.3389/fimmu.2022.808607

55. Dubensky TW, Kanne DB, Leong ML. Rationale, progress and development of vaccines utilizing STING-activating cyclic dinucleotide adjuvants. Ther Adv Vaccines. 2013 Nov;1(4):131–43.

56. Li XD, Wu J, Gao D, Wang H, Sun L, Chen ZJ. Pivotal roles of cGAS-cGAMP signaling in antiviral defense and immune adjuvant effects. Science. 2013 Sep 20;341(6152):1390–4.

57. Khoo LT, Chen L. Role of the cGAS–STING pathway in cancer development and oncotherapeutic approaches. EMBO Rep. 2018 Dec;19(12):e46935.

58. Michael Lavigne G, Russell H, Sherry B, Ke R. Autocrine and paracrine interferon signalling as ‘ring vaccination’ and ‘contact tracing’ strategies to suppress virus infection in a host. Proc Biol Sci. 288(1945):20203002.

59. Reed LJ, Muench H. A SIMPLE METHOD OF ESTIMATING FIFTY PER CENT ENDPOINTS12. American Journal of Epidemiology. 1938 May;27(3):493–7.

